# Gp78 E3 ubiquitin ligase mediates both basal and damage-induced mitophagy

**DOI:** 10.1101/407593

**Authors:** Bharat Joshi, Yayha Mohammadzadeh, Guang Gao, Ivan R. Nabi

## Abstract

Mitophagy, the elimination of mitochondria by the autophagy machinery, evolved to monitor mitochondrial health and maintain mitochondrial integrity. PINK1 is a sensor of mitochondrial health that recruits Parkin and other mitophagy-inducing ubiquitin ligases to depolarized mitochondria. However, mechanisms underlying mitophagic control of mitochondrial homeostasis, basal mitophagy, remain poorly understood. The Gp78 E3 ubiquitin ligase, an endoplasmic reticulum membrane protein, induces mitochondrial fission, endoplasmic reticulum-mitochondria contacts and mitophagy of depolarized mitochondria. CRISPR/Cas9 knockout of Gp78 in HT-1080 fibrosarcoma cells results in reduced ER-mitochondria contacts, increased mitochondrial volume and resistance to CCCP-induced mitophagy. Knockdown (KD) of the essential autophagy protein ATG5 increased mitochondrial volume of wild-type cells but did not impact mitochondrial volume of Gp78 knockout cells. This suggests that endogenous Gp78 actively eliminates mitochondria by autophagy in wild-type HT-1080 cells. Damage-induced mitophagy of depolarized mitochondria, in the presence of CCCP, but not basal mitophagy was prevented by knockdown of PINK1. This suggests that endogenous Gp78 plays dual roles in mitophagy induction: 1) control of mitochondrial homeostasis through mitophagy of undamaged mitochondria; and 2) elimination of damaged mitochondria through PINK1.

## Introduction

Mitophagy, mitochondrial autophagy, is a process by which mitochondria are selectively targeted for autophagy and lysosomal degradation (1–3). Mitophagy plays a critical role in cellular health by controlling mitochondrial mass and eliminating damaged or dysfunctional mitochondria (4). A number of molecular pathways have been described that mediate mitophagy, including the best-characterised PINK1/Parkin pathway, closely associated with Parkinsons disease (5). In response to mitochondrial damage, loss of mitochondrial inner membrane potential prevents removal of PTEN-induced putative kinase 1 (PINK1) that accumulates on the outer mitochondrial membrane (OMM) (6). PINK1 phosphorylates pre-existing ubiquitin on OMM proteins, recruits the E3 ligase Parkin to damaged mitochondria and phosphorylates Parkin activating its ubiquitin ligase activity. Activated Parkin adds ubiquitin to OMM proteins which can again be phosphorylated by PINK1 amplifying a feedforward cycle of Parkin recruitment and activation by phosphorylating Parkin and ubiquitin (3). Parkin ubiquitination of OMM proteins of damaged mitochondria triggers autophagosome formation through recruitment of LC3 and the autophagy machinery (7).

However, while the mechanism of action of PINK1/Parkin in mitophagy has been extensively characterized, the majority of studies have been based on Parkin overexpression and mitochondrial depolarization with CCCP (2, 6, 8–10). Further, many cell lines do not express Parkin, tissue expression of Parkin is varied and mitophagy independent of Parkin and/or PINK1 has been reported (11–13). Ubiquitin and PINK1/Parkin-independent mitophagy occurs through receptor-mediated pathways that include the BNIP3/NIX/FUNDC1-induced mitophagy in response to hypoxia (2, 14). Other ubiquitin ligases such as Mul1, MARCH, SMURF, and Gp78 have been reported to function independently or in parallel with Parkin (10, 15–18). PINK1-dependent, Parkin-independent mitophagy pathways include the synphilin/SIAH1 and the Mulan pathways (15, 16, 19). However, synphilin is primarily expressed in brain and whether Mulan functions downstream of or independently of PINK1 remains to be determined (1). Nevertheless, PINK1 remains, to date, a unique sensor of mitochondrial damage (20, 21). A major challenge to understanding mitophagy regulation is the demonstration of an ubiquitin ligase, other than Parkin, that functions endogenously to control basal mitochondrial homeostasis and mediate PINK1-dependent mitophagy of damaged mitochondria.

Gp78 is a key E3 ubiquitin ligase in the ER-associated degradation (ERAD) pathway that targets misfolded and physiological substrates for ubiquitination and degradation by the proteasome (22–27). Gp78 ubiquitin ligase activity promotes ER-mitochondria contacts and targets the mitofusins for degradation inducing mitochondrial fission and enhances mitophagy upon mitochondrial depolarization (17, 28, 29). We show here that CRISPR/Cas9 Gp78 knockout results in increased mitochondrial volume and prevents CCCP-induced mitophagy, defining a role for endogenous Gp78 in regulation of both basal and damage-induced mitophagy. We further show that Gp78-dependent mitophagy of damaged mitochondria is PINK1-dependent.

## Materials and Methods

### Cell lines, antibodies and reagents

The HT-1080 fibrosarcoma cell line was acquired from ATCC, authenticated by STR profiling (TCAG Genetic Analysis Facility, Hospital for Sick Kids, Toronto, ON, Canada, www.tcag.ca/facilities/geneticAnalysis.html), tested regularly for mycoplasma infection by PCR (ABM, Richmond, BC, Canada) and maintained in RPMI 1604 media supplemented with 10% FBS and 1% L-Glutamine in a 37°C incubator with 5% CO_2_. HT-1080 and Gp78 CRISPR clones were allowed to grow only up to six passages. Cells were passed every 48 hours at a density of 300,000 cells per 10 cm petri dish, rinsed every 24 hours with 10 ml of PBS and supplied with 10ml of fresh complete medium.

Anti-ATPB pAb/mAb, anti-MFN1 mAb (ab128743/ab5432 and ab57602) and anti PINK1 pAb (ab23703), were purchased from Abcam (Cambridge, MA, USA), anti-Gp78 pAb (#16675-1-AP) from Proteintech (USA) and anti-MFN2 pAb (sc-50331) from SantaCruz Biotech (USA). Anti-STX17 pAb (HPA001204), anti-β-Actin mAb (A5441), tissue culture grade DMSO (D2650), and CCCP (C2759) were purchased from Sigma (USA). EGFP-LC3 was from Karla Kirkegaard (Addgene plasmid #11546) (30). YFP-Parkin was from Richard Youle (Addgene plasmid #23955) (8). Flag-Gp78 was as previously described (31).

### CRISPR/Cas9 knockout of Gp78

GeneArt-Crispr/Cas9 Nuclease vector with OFP (Orange Fluorescence Protein) kit (A21174, Invitrogen, USA) was from Lifetechnologies, Thermo Fisher. We used http://crispr.mit.edu to design guided RNAs and http://www.rgenome.net/cas-offinder/ (RGEN tools) to check them for off-target effects and used the following oligonucleotides for guide RNA1 (5’-CAC CGG AGG AAG AGC AGC GGC ATG G-3’, 5’-AAA CCC ATG CCG CTG CTC TTC CTC C-3’) and guide RNA2 (5’-CAC CGG CCC AGC CTC CGC ACC TAC A-3’, 5’-AAA CTG TAG GTG CGG AGG CTG GGC C-3’). The guide RNAs were in vitro annealed, cloned into the GeneArt linear vector according to the supplier’s protocol and sequence verified prior to transfection into HT-1080 cells. Sequence verified gRNA1 or gRNA2 containing GeneArt-Crispr/Cas9 Nuclease vector with OFP were transiently transfected into HT-1080 cells, plated 24 hours previously, using Effectene transfection reagent (Cat. #301425, Qiagen, USA). After 36 hours incubation cells were harvested and genomic DNA was isolated to perform GeneArt Genomic Cleavage Detection assay (A24372, Invitrogen, USA) to check cleavage efficiency. Once cleavage efficiency was confirmed, HT-1080 cells were replated for 36 hours, trypsinized, FACS sorted and OFP expressing cells were singly plated in 96 well pates by serial dilution. Single colonies were replicated in 12 well plates and a set was frozen and stored in liqN2 whereas the other set was subjected to lysate preparation, SDS PAGE and Gp78 western blot analysis. Arbitrarily chosen representative clones (g1-3, g1-4, g1-7; g2-13, g2-36, g2-41) from both gRNAs were expanded, tested for mycoplasma and stored as multiple freezedowns. From isolated genomic DNA, an approximate 800bp fragment flanking Exon1 of Gp78 was PCR amplified using Q5 (Qiagen, USA) the following primer set (Forward: 5’-CTG GAG GCT ACT AGC AAA-3’, Reverse: 5’-ATG TGG CCC AGT ACC T-3’) and TA cloned. At least ten clones were sequenced from each to confirm INDEL.

### siRNA knockdown, plasmid transfection and western blotting

siControl, siPINK1 and siATG5 (#D-001810-01-05, #L-004030-00-0005, #L-004374-00-0005) were purchased from Dharmacon and transiently transfected wherever indicated to wild-type HT-1080 cells or g1-4 or g2-41 Gp78 CRISPR clones using Lipofectamine 2000 (#11668019, Invitrogen, USA) following the manufacturer’s protocol. All siRNA transfection experiments were for 48 hours and treatments were performed 24 hours post siRNA transfection. Alternatively, cells were transiently transfected with mammalian protein expressing plasmids (pcDNA3, LC3-GFP, Flag-Gp78, YFP-Parkin) using Effectene (Qiagen, Germany) following the manufacturer’s protocol. Where indicated, cells were treated with 10 μM of CCCP or a corresponding volume of DMSO as control 24 hours prior to fixation or harvesting cells. Western blotting was performed as previously described using Horseradish Peroxidase (HRP)-conjugated secondary antibody followed by addition of ECL (GE Healthcare Bio-Sciences Corp., USA) to reveal chemiluminescence (32). Densitometry quantification was done using ImageJ software.

### Immunofluorescent labelling and confocal microscopy

For immunofluorescent labeling, cells were: (1) fixed with 3.0% PFA for 15 minutes at room temperature and washed with PBS-CM (phosphate buffer solution supplemented with 1 mM CaCl2 and 10 mM MgCl2); (2) permeabilized with 0.2% Triton X-100 and washed with PBS-CM; (3) blocked with 1% BSA for 1 hour at room temperature; (4) labeled with anti-ATPB for one hour followed by washing with PBS-CM; (5) incubated with secondary antibodies for 1 hour followed by washing with PBS-CM; and (6) mounted in ProLong Diamond (Thermo Fisher) and cured for 24 hours before imaging. Confocal image stacks were obtained on a III-Zeiss spinning disk confocal microscope (Intelligent Imaging Innovations Inc.) with either Zeiss Plan-Apochromat 63X/1.4NA or 100X/1.4NA oil objectives using Slidebook image acquisition and analysis software (LSI Imaging Facility, Life Sciences Institute, UBC). ATPB mitochondrial labeling was thresholded from 3D images to measure mitochondrial volume. To measure the number of the LC3 puncta associated with mitochondria, the LC3-GFP image was thresholded to select only for puncta and not nuclear or dispersed cytoplasmic signal and the number of LC3 puncta that overlapped with the mitochondrial mask were counted.

## Results

### Gp78 CRISPR/Cas9 knockout HT-1080 cells

The HT-1080 fibrosarcoma cell line expresses high levels of Gp78 protein and has been extensively used for the study of Gp78 (28, 33, 34). We previously used stable miRNA and inducible lenti-shRNA approaches to post-transcriptionally knockdown Gp78 mRNA (17, 28, 35, 36). We now applied the CRISPR/Cas9 technique to knockout Gp78 using two different guided RNA sequences: gRNA1 that targets and eliminates the ATG start codon; and gRNA2 that induces a frameshift sixteen amino acids downstream of the ATG start codon (Figure 1A). Complementary gRNA1 or gRNA2 oligos were annealed and cloned in GeneArt-OFP plasmid, sequence verified and transfected into HT-1080 cells. We screened sixty clones by Western blotting and obtained 38 Gp78 knockout clones. Three clones showing complete absence of Gp78 from each gRNA were chosen (gRNA1: clones #3, 4 and 7; gRNA2: clones #13, 36 and 41) for further experimentation. From isolated genomic DNA for each clone, the DNA fragment flanking Exon1 of Gp78 was PCR amplified, cloned and sequenced. As shown in Figure 1A, for gRNA1 clones #3 and #4 the G was deleted from the ATG start codon while for clone #7 an additional T was inserted in the start codon generating ATTG. For all three gRNA2 clones (#13, 36, 41), a T was inserted at amino acid 16 (CCTA to CCTTA) causing a frameshift mutation. As shown in Figure 1A, probing Western blots with anti-Gp78 polyclonal antibody shows the complete absence of Gp78 in all six gRNA1 and gRNA2 clones compared to wild-type HT-1080 cells. We previously reported that Gp78 shRNA knockdown was associated with increased ER-mitochondria interaction (28) using Syntaxin 17 (STX17), a MAM localized protein, as a reporter (37). Similarly, 3D spinning disk confocal analysis of Gp78 CRISPR/Cas9 knockout clone 41 shows that individual mitochondria were larger and had reduced overlap with STX17 compared to wild-type HT-1080 cells (Figure 1B).

**Figure 1:**
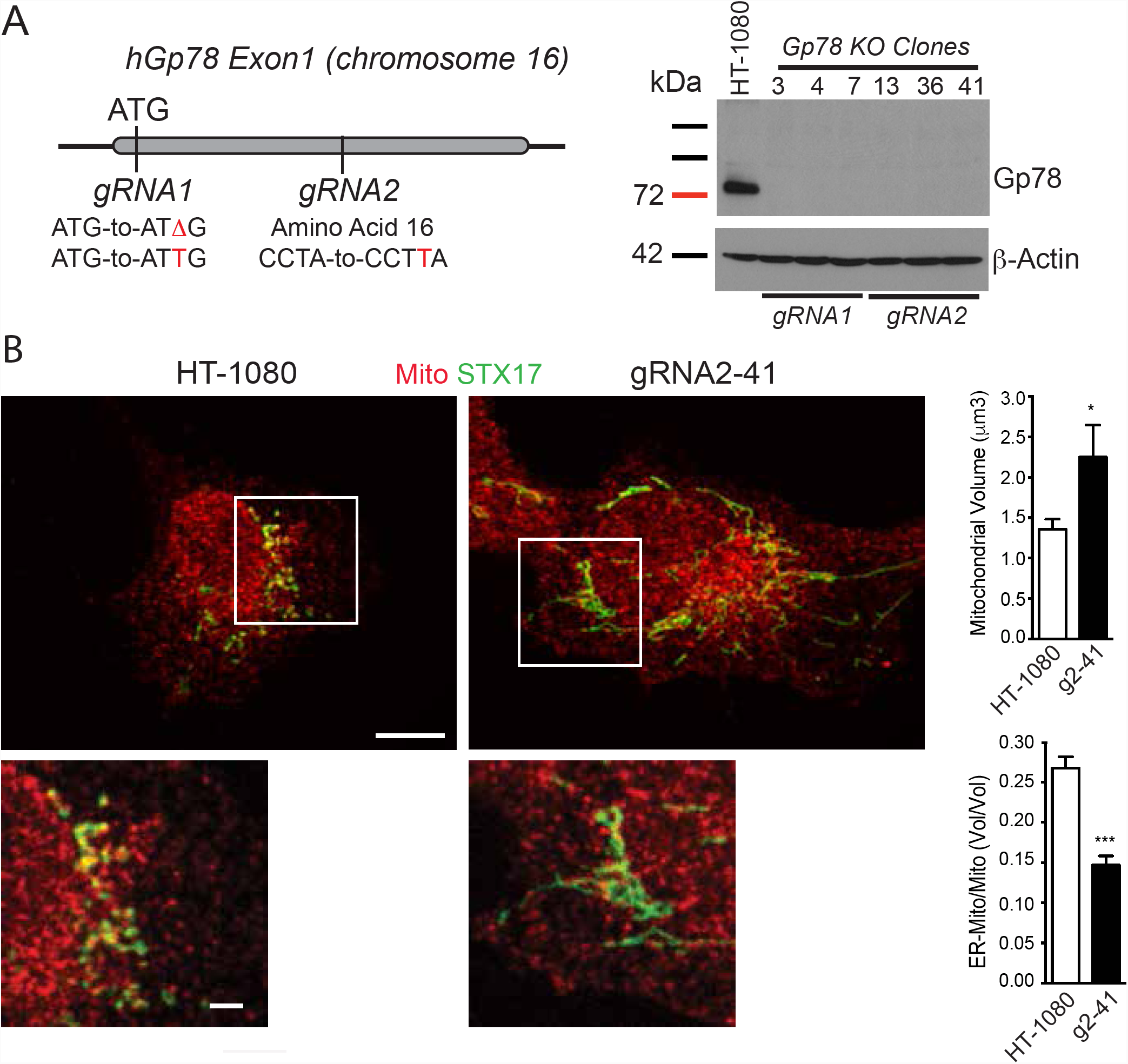
Generation of Gp78 knockout HT-1080 cells using CRISPR/Cas9. **A)** Schematic showing Exon 1 region of Gp78 gene on chromosome 16 and location of gRNA1 targeting the start codon, deleting ATG (δG or inserting extra T), and of gRNA2, inserting an extra T at amino acid 16 causing frameshift and termination. Western blot for Gp78 is shown for wild-type HT-1080 cells and all six gRNA1 and gRNA2 Gp78 knockout CRISPR clones with β-actin as a loading control. **B)** 3D spinning disk confocal microscopy of HT-1080 and CRISPR clone #g2-41 labeled for the mitochondrial protein ATPB and the ER-mitochondria marker STX17. Quantification show significantly increase volume of individual mitochondrial and reduced STX17-mitochondria overlap (n=3 or >3, *, p < 0.05; ***, p<0.001; ±SEM; Scale Bars: 5 μm; 1 μm for zooms).

### Gp78 knockout increases mitochondrial volume, reduces ER-mitochondria contacts and limits mitophagy in HT-1080 cells

Gp78 CRISPR KO HT-1080 cells labeled for the inner mitochondrial protein ATBP all show extended mitochondrial networks compared to wild-type HT-1080 cells. Quantification of total mitochondrial volume from 3D spinning disk confocal stacks shows a significant ∼2-fold increase in mitochondrial volume in the Gp78 knockout clones relative to wild-type HT-1080 cells (Figure 2). To determine if Gp78 knockout specifically affected mitophagy, we assessed the impact of the mitochondrial membrane potential decoupler, CCCP (10 μM; 24h) on mitochondrial volume in wild-type and Gp78 CRISPR KO HT-1080 cells. CCCP treatment induces a significant reduction in mitochondrial volume in wild-type HT-1080 but does not impact the elevated mitochondrial volume of the Gp78 KO clones (Figure 2). This indicates that endogenous Gp78 is required for mitophagy in HT-1080 cells.

**Figure 2:**
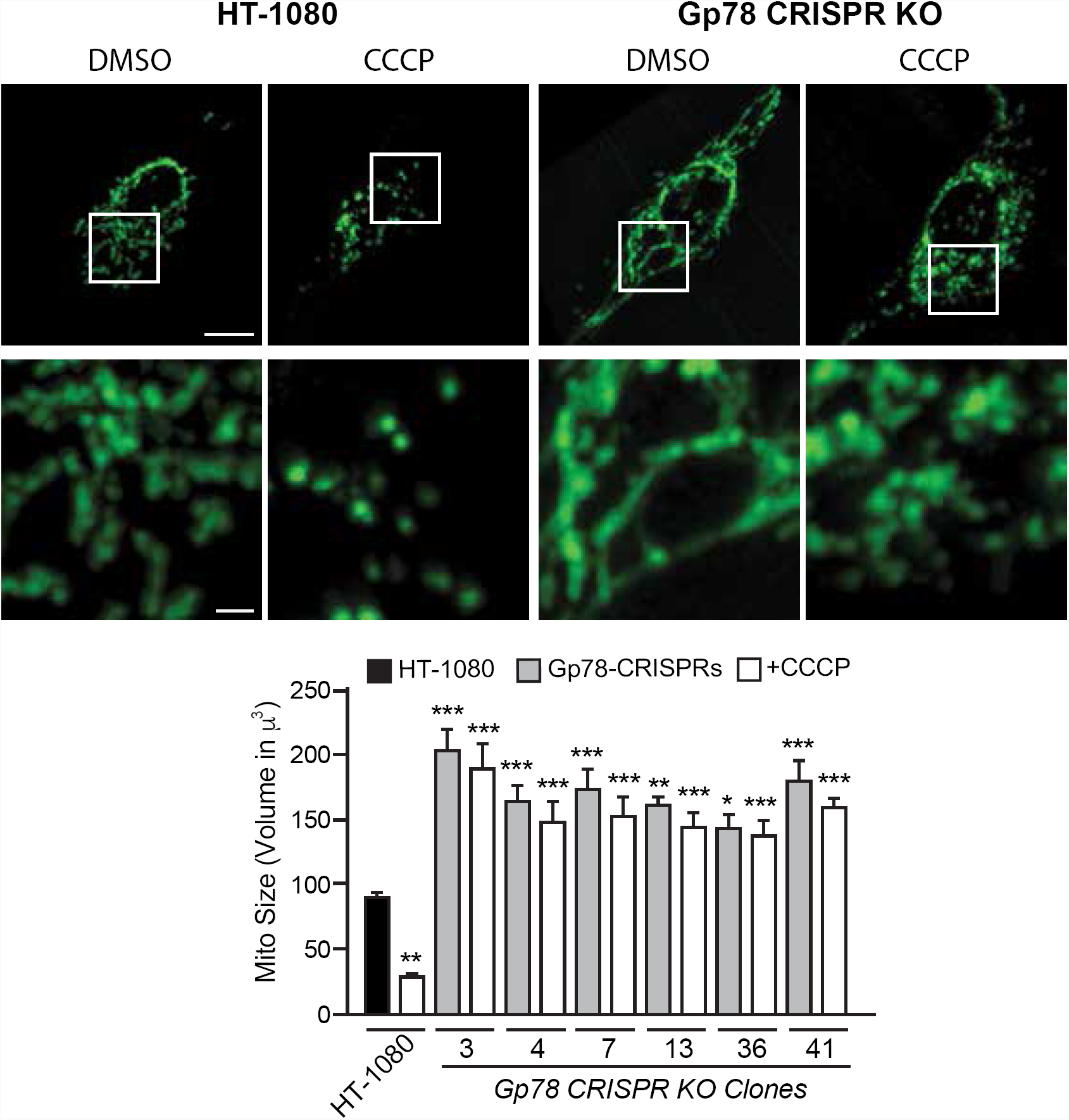
Gp78 CRISPR/Cas9 knockout prevents damage-induced mitophagy. Wild-type HT-1080 cells and the six Gp78 knockout CRISPR clones were incubated with either DMSO or 10 μM CCCP for 24 hours then fixed and labeled for mitochondrial ATPB and imaged by 3D spinning disk confocal microscopy. Representative images of HT-1080 and a CRISPR clone including magnification of boxed region are shown. Quantification of total mitochondrial volume is shown in the bar graph (n=3; *, p<0.05; **, p<0.01; ***, p<0.001 relative to DMSO or CCCP treated HT-1080 cells, respectively; ±SEM; Scale Bars: 5 μm; 1 μm for zooms).

Consistent with the increased mitochondrial volume of Gp78 CRISPR KO HT-1080 cells, by Western blot analysis ATPB levels show a significant two-fold elevation in Gp78 CRISPR clones relative to wild-type HT-1080 cells (Figure 3A). To test whether basal Gp78-dependent mitophagy was responsible for the reduced mitochondria levels of Gp78 CRSPR clones, we knocked down the essential autophagy gene ATG5. ATG5 siRNA knockdown increases ATPB levels in HT-1080 cells to those of Gp78 KO CRISPR cells (Figure 3A). Similarly, ATG5 knockdown increases mitochondrial volume by 3D spinning disk confocal analysis of both untreated and CCCP-treated wild-type HT-1080 cells (Figure 3B). ATG5 knockdown does not affect mitochondrial volume in the Gp78 KO clones in the absence or presence of CCCP. This suggests that the increased basal mitochondrial volume of Gp78 knockout cells, in the absence of CCCP, is due to inhibition of active Gp78-dependent mitophagy in wild-type HT-1080 cells.

**Figure 3:**
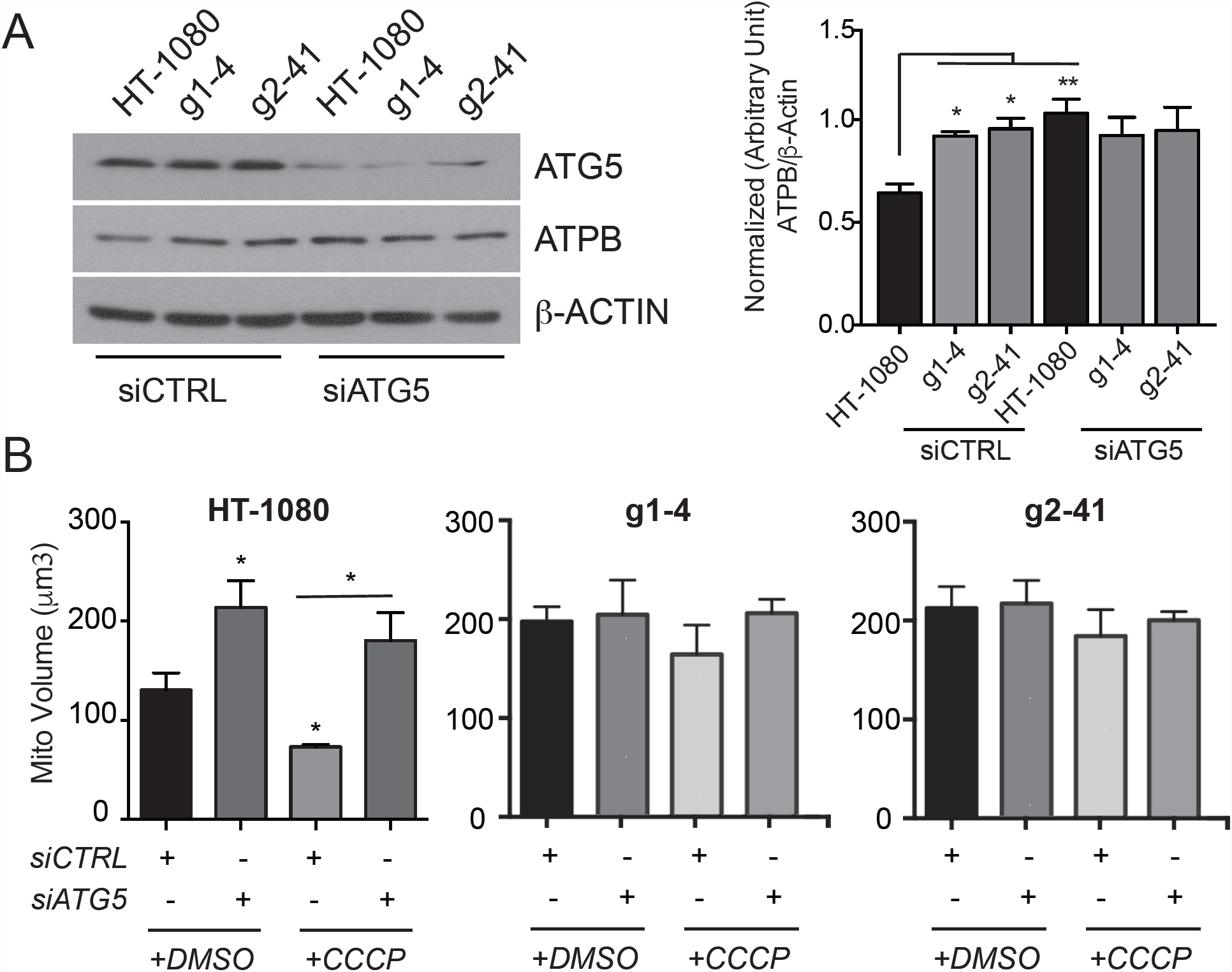
Gp78 induces basal mitophagy in HT-1080 cells. **A)** Wild-type HT-1080 cells and the g1-4 and g2-41 Gp78 knockout CRISPR clones were transfected with non-specific siCTL or siATG5 and western blotted for ATG5, mitochondrial ATPB and β-actin. Densitometry quantification of ATBP relative to β-actin is shown in the bar graph. **B)** Wild-type HT-1080 cells and the g1-4 and g2-41 Gp78 knockout CRISPR clones transfected with non-specific siCTL or siATG5 were incubated with either DMSO or 10 μM CCCP for 24 hours and then fixed and labeled for mitochondrial ATPB and imaged by 3D spinning disk confocal microscopy. Quantification of total mitochondrial volume is shown in the bar graph. (n=3, *, p < 0.05; **, p<0.01; ±SEM).

### Gp78 mitophagy of damaged mitochondria is PINK1-dependent

The Parkin, SIAH1 and Mulan ubiquitin ligases (15–17) function downstream of PINK1 to promote mitophagy of damaged mitochondria. HT-1080 cells do not express Parkin and Gp78 induces mitophagy independently of Parkin (17). We therefore tested whether PINK1 mediated Gp78-dependent mitophagy. As seen in Figure 4 A,B, siRNA knockdown of PINK1 prevented the CCCP-induced reduction of mitochondrial volume in wild-type HT-1080 cells. Mitochondria in CCCP-treated siPINK1 cells were fragmented but retained the total volume of untreated HT-1080 cells. Unlike ATG5 siRNA, PINK1 siRNA did not affect mitochondrial volume of HT-1080 wild-type cells in the absence of mitochondrial damage (Figure 4A,B). This indicates that PINK1 selectively mediates Gp78-dependent mitophagy of damaged mitochondria and not the basal Gp78-dependent mitophagy in HT-1080 cells. To confirm that PINK1 was modulating mitochondrial volume through mitophagy, we expressed EGFP-LC3 in HT-1080 wild-type and knockout cell lines and assessed the overlap between LC3 puncta and mitochondria in response to CCCP (Figure 4A,C). CCCP induced a significant increase in the number of LC3 puncta associated with mitochondria in WT HT-1080 cells compared to Gp78 knockout cells. However, upon CCCP treatment, the number of mitochondrial associated LC3 puncta was reduced in siPINK1 transfected wild-type HT-1080 cells to the same level as the Gp78 CRISPR/Cas9 knockout cell lines. This indicates that mitophagy of damaged mitochondria by endogenous Gp78 depends on PINK1.

**Figure 4:**
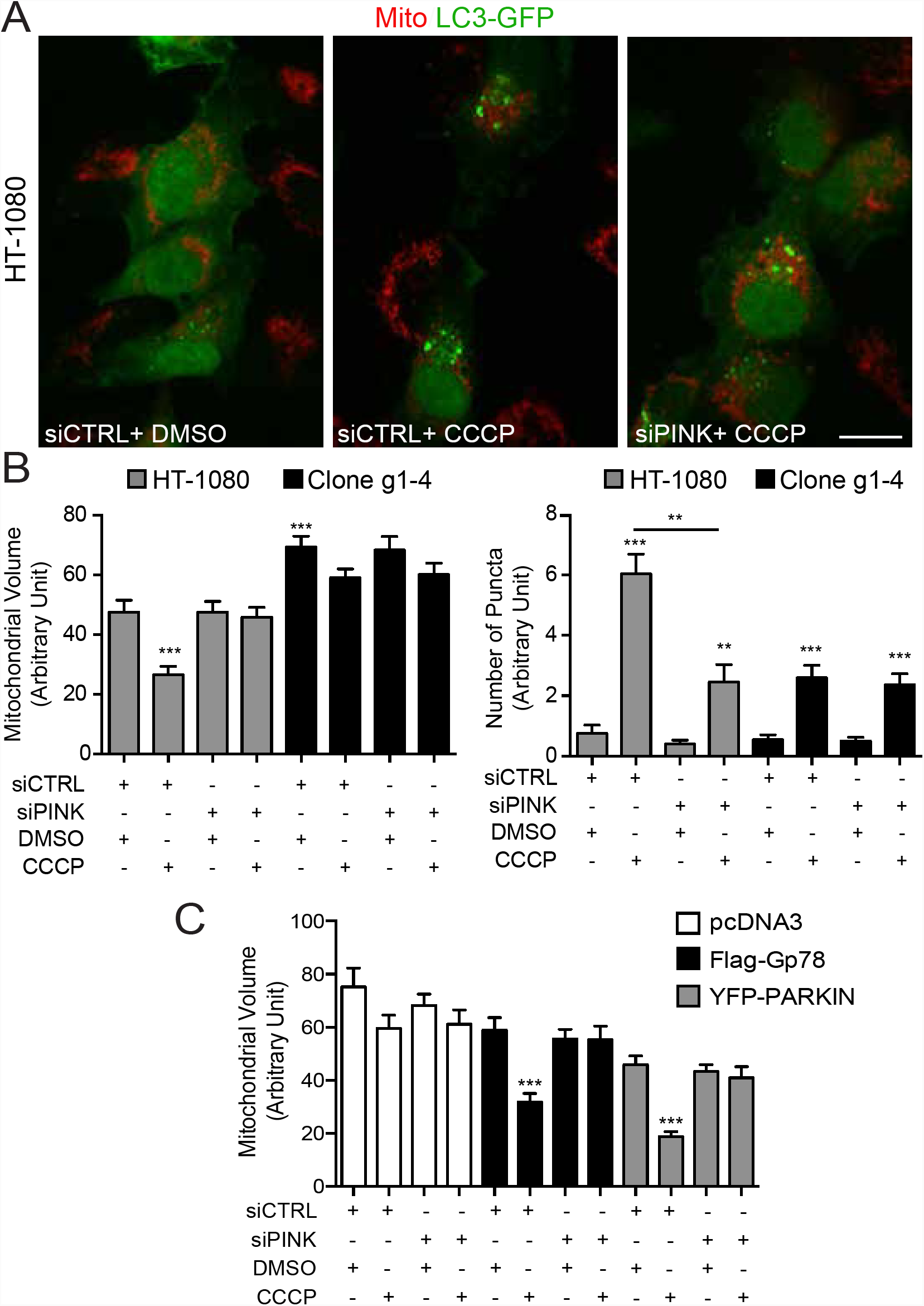
Gp78 mitophagy of damaged mitochondria is PINK1-dependent. **A)** Wild-type HT-1080 and Gp78 CRISPR g1-4 cells were transiently transfected with LC3-GFP and either siCTL or siPINK1, incubated with either DMSO or 10 μM CCCP for 24 hours and then fixed and labeled for mitochondrial ATPB and imaged by 3D spinning disk confocal microscopy. Representative ATPB labeled mitochondria are shown in red and LC3-GFP labeling in green. **B)** Quantification of total mitochondrial volume (left) and LC3-puncta overlap with mitochondria determined by mask overlay analysis (right) are shown in the bar graphs. (n=3, *, p < 0.05; **, p<0.01; ±SEM). **C)** Gp78 CRISPR g1-4 clone was transfected with pcDNA, Flag-Gp78 or YFP-Parkin and either siCTL or siPINK1, incubated with either DMSO or 10 μM CCCP for 24 hours and then fixed and labeled for mitochondrial ATPB and imaged by 3D spinning disk confocal microscopy. Quantification of total mitochondrial volume is shown in the bar graph. (n=3, **, p < 0.01; ***, p<0.001; ±SEM; Scale Bar: 20 μm).

Gp78 induces mitophagy in cells devoid of Parkin (37). To test whether Parkin mitophagy is independent of Gp78, we overexpressed Parkin in HT-1080 CRISPR/Cas9 Gp78 KO cells and treated the cells with siPINK1. Upon induction of mitochondrial damage with CCCP, YFP-Parkin overexpression induced PINK1-dependent mitophagy in the absence of endogenous Gp78. In addition, Flag-Gp78 overexpression rescued mitophagy in the HT-1080 KO cell line in a PINK1-dependent manner (Figure 4C). This suggests that Gp78 and Parkin act independently downstream of PINK1 to promote mitophagy. PINK1 identification of damaged mitochondria therefore represents, at least to date, a unique cellular mechanism for the identification of damaged mitochondria to be targeted for degradation. That diverse ubiquitin ligases independently mediate PINK1-dependent mitophagy, as shown here for Gp78 and Parkin, highlights the multiple and likely cell-specific pathways that are responsible for this critical cellular process.

Importantly, we show here that Gp78 promotes basal mitophagy in the absence of mitochondrial damage in a PINK1-independent manner. Knockdown of the essential autophagy gene ATG5 increases mitochondrial volume in wild-type HT-1080 cells to levels of Gp78 knockout cells, defining a critical role for Gp78 control of basal mitophagy in the homeostatic regulation of mitochondrial levels independently of mitochondrial damage. PINK1 knockdown restored mitochondrial volume to levels of wild-type HT-1080 cells, and not that of HT-1080 Gp78 knockout cells suggesting that distinct mechanisms are responsible for Gp78-dependent basal mitophagy and mitophagy of damaged mitochondria. Indeed, our result is in line with a recent study of PINK1 knockout mice showing that PINK1 is dispensable for basal mitophagy *in vivo* and stating that the proteins involved in basal mitophagy are hitherto unknown (12). Gp78 is a key E3 ubiquitin ligase in ER-associated degradation (22, 27) and the demonstration here that it mediates basal mitophagy highlights a molecular connection between these two processes.

Gp78 promotes the formation of rough ER-mitochondria contacts and recruits Syntaxin17 to mitochondria, consistent with the established role of Syntaxin17 in mitophagy (28, 37). The ability of Parkin to induce CCCP-dependent mitophagy in Gp78 knockout cells, that show a reduction in Syntaxin 17-mitochondria contacts, suggests that Gp78 induction of ER-mitochondria contacts is not required for Parkin-induced mitophagy of damaged mitochondria. This is consistent with a recent study that argues that disruption of ER-mitochondria contacts is associated with Parkin-mediated mitophagy (38). This study highlights the diverse mechanisms that promote mitophagy and the specific role of Gp78-dependent rough ER-mitochondria contacts in mitophagy remains to be determined.

## Acknowledgements

This study was supported by a grant from the Canadian Institutes of Health Research)CIHR Grant PJT-148698) and 1Y and 4Y UBC Fellowships (YM, GG).

## Supp. Video

Mitochondrial ATPB (red) and LC3-GFP in a CCCP-treated HT-1080 cell.

